# MAFFIN: Metabolomics Sample Normalization Using Maximal Density Fold Change with High-Quality Metabolic Features and Corrected Signal Intensities

**DOI:** 10.1101/2021.12.23.474041

**Authors:** Huaxu Yu, Tao Huan

**Affiliations:** Department of Chemistry, Faculty of Science, University of British Columbia, Vancouver Campus, 2036 Main Mall, Vancouver, V6T 1Z1, BC, Canada

## Abstract

Sample normalization is a critical step in metabolomics to remove differences in total sample amount or concentration of metabolites between biological samples. Here, we present MAFFIN, an accurate and robust post-acquisition sample normalization workflow that works universally for metabolomics data collected by mass spectrometry (MS)-based platforms. The most important design of MAFFIN is the calculation of normalization factor using maximal density fold change (MDFC) value computed by a kernel density-based approach. MDFC is more accurate than traditional median FC-based normalization, especially when the numbers of up- and down-regulated metabolic features are different. In addition, we showcase two essential steps that are overlooked by conventional normalization methods, and incorporated them into MAFFIN. First, instead of using all detected metabolic features, MAFFIN automatically extracts and uses only the high-quality features to calculate FCs and determine the normalization factor. In particular, multiple orthogonal criteria are proposed to pick up the high-quality features. Second, to guarantee the accuracy of the FCs, the MS signal intensities of the high-quality features are corrected using serial quality control (QC) samples. Using simulated data and urine metabolomics datasets, we demonstrated the critical need of high-quality feature selection, MS signal correction, and MDFC. We also show the superior performance of MAFFIN over other commonly used post-acquisition sample normalization methods. Finally, a biological application on a human saliva metabolomics study shows that MAFFIN provides robust sample normalization, leading to better data separation in principal component analysis (PCA) and the identification of more significantly altered metabolic features.

**TOC:** 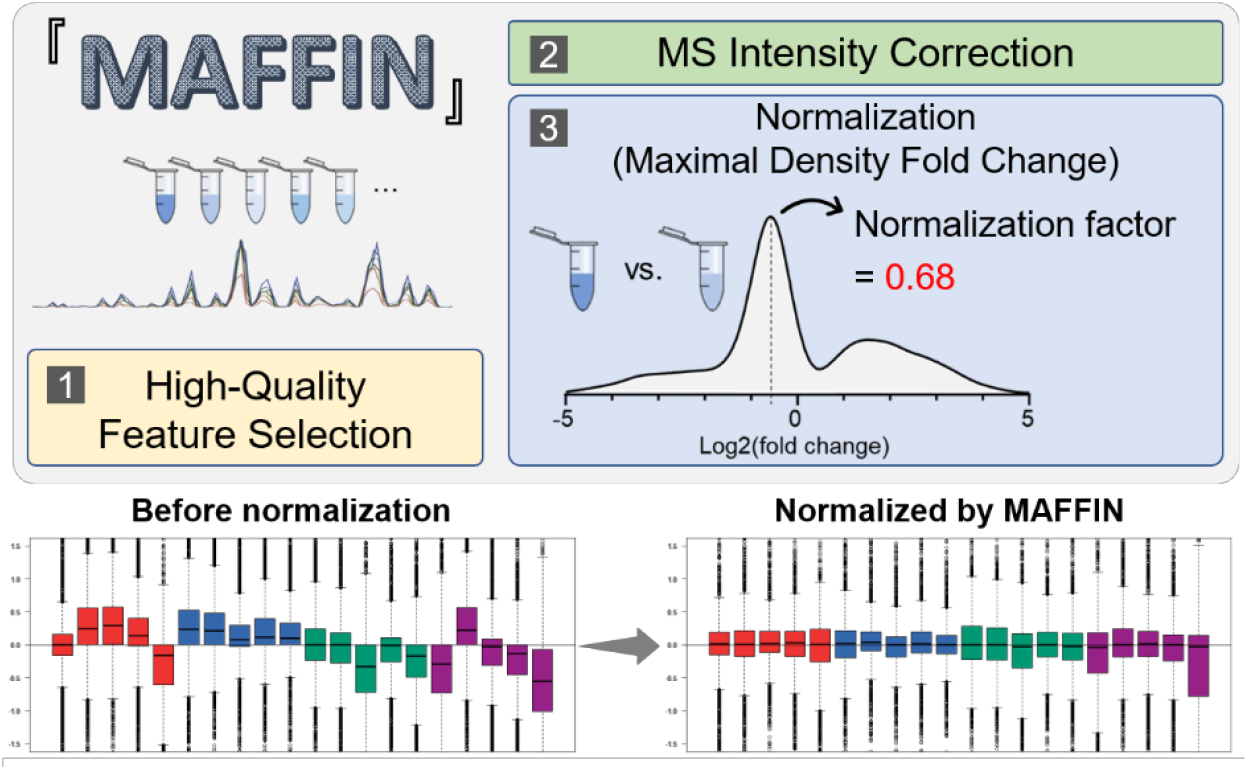

## Introduction

Liquid chromatography-mass spectrometry (LC-MS)-based untargeted metabolomics enables high-throughput and high-coverage detection and quantification of thousands of metabolic features in various biological sample types.^1–3^ However, many biological samples types, such as urine, saliva, sweat, and fecal, have large differences in total sample amount or concentration of metabolites. For instance, urine samples can have a greater than 15-fold concentration difference because of water intake or other physiological factors.^4^ For human fecal samples, its water content can commonly change from 60% to 85% even within a week.^5, 6^ Besides this biology-induced dilution effect, inappropriate analytical procedure can also induce significant sample-to-sample variation. For example, the mass of a small amount of frozen tissue (e.g., less than 1 mg) can be significantly affected by the moisture absorbed from air during the weighing process.^7^ Likewise, cell counts performed before scraping cells off of a petri dish can be inaccurate if some cells are left behind.^8^ These analytical and biological variations cause different total sample amount between comparative groups, leading to larger variation and/or systematic biases in quantitative metabolomics comparisons.

The metabolomics community has recognized the importance of sample normalization. Various normalization strategies have been proposed to assure accurate and precise quantitative comparison and statistical analysis. These normalization methods can usually be classified into pre- or post-acquisition sample normalization.^9^ Pre-acquisition sample normalization is applied prior to MS data collection. It measures a certain quantity that reflects the total sample amount or concentrations of metabolites. The samples are then reconstituted to appropriate final volumes based on the measured quantities to make the total concentration consistent between the samples.

For example, creatinine levels generally reflect the urine concentration and is commonly used for normalizing urine samples.^4, 10, 11^ For cell samples, total DNA or protein concentration is expected to agree with the quantity of cells and thus are often used to normalize cell samples when cell counts are not accurate.^8^ Recent work has used dansylation labeling to quantify the total amount of amine- and phenol-containing metabolites for sample normalization.^12, 13^ The underlying assumption of that work is that the total amount of amine- and phenol-containing metabolites is a good reflection of the total metabolome, and it should be consistent in a biological sample type that is involved in a metabolomics study. In general, pre-acquisition sample normalization is often favored over post-acquisition sample normalization. One key benefit of pre-acquisition sample normalization is that researchers can estimate the total amount of metabolites and then optimize sample loading amount on the LC-MS system for better analytical performance.^14^ In addition, pre-acquisition sample normalization also enables comparisons across different sample batches and sample quality checks. As such, pre-acquisition sample normalization is preferred if a proper quantity that indicates the total metabolome concentration is available. Unfortunately, for many biological sample types, such quantity either does not always work or does not exist. For instance, the urine creatinine normalization was reported to have bias, especially when glomerular filtration rate is changed due to kidney injury^10^. On the other side, many samples types, including but not limited to saliva, sebum, and sweat, still lack reliable quantities for pre-acquisition sample normalization.

Post-acquisition sample normalization is an alternative strategy that complements and addresses the limitations of pre-acquisition sample normalization. It is a data driven approach that adjusts the MS signal intensities of the collected data based on a certain criterion that is considered to be unchanged across samples. Commonly used post-acquisition normalization methods include the use of mass spectrum total useful signal (MSTUS) or total useful peak area (TUPA), quantile normalization, probabilistic quotient normalization (PQN), and median fold change.^15–19^ However, these methods perform inadequately due to their inherent limitations.^9^ In particular, problem with total peak intensity, problem with medium fold change. In this work, we developed MAFFIN, a universal post-acquisition normalization method that is adaptable to any sample type. Using both simulated and real metabolomics data sets, we demonstrated that the MAFFIN workflow outperforms four other normalization methods, including sum normalization, median normalization, probabilistic quotient normalization, and quantile normalization, in reducing the biological variation. The normalized data can largely improve the confidence of downstream statistical analysis and biological interpretations.

## Methods

### Chemicals

Methanol (MeOH), water (H2O), acetonitrile (ACN), sodium formate (NaFA), and ammonium acetate (all LC-MS-grade) were purchased from Thermo Fisher Scientific (Waltham, MA). Ammonium hydroxide was obtained from MilliporeSigma (Burlington, MA).

### Urine and Saliva Sample Preparation

The human urine and saliva samples were collected from four healthy volunteers, including 2 males and 2 females. For each participant, 5 samples were collected at random time points to mimic different sample concentrations. The collected biospecimens were stored at −80 °C prior to the metabolomics experiment. For metabolome extraction, 100 μL of urine was mixed with 1 mL of ice-cold methanol in a 1.5 mL Eppendorf vial and vortexed. The solution was incubated at −20 °C for 2 hrs, followed by centrifugation at 14,000 rpm for 15 min at 4 °C. The clear supernatant was dried in a SpeedVac at 20 °C and then reconstituted in 150 μL ACN and H2O (1:1, v:v) mixed solvent for LC-MS analysis. The method blank was also prepared following the same protocol but without urine. The QC sample was prepared by pooling 20 μL aliquots from each sample. The saliva samples were prepared following the same workflow.

### LC-MS Analysis

LC-MS analysis was performed on a UHR-QqTOF (ultra-high resolution Qq-time-of-flight) mass spectrometer Impact II (Bruker Daltonics, Bremen, Germany) coupled with a 1290 Infinity II LC system (Agilent Technologies, Palo Alto, CA). Hydrophilic interaction chromatography (HILIC) separation was performed on a SeQuant ZIC-pHILIC column (150 mm × 2.1 mm, 5 μm, 200 Å). Mobile phase A was 5% ACN in H2O with 10 mM ammonium acetate (pH = 9.8, adjusted by ammonium hydroxide), and mobile phase B was 5% H2O in ACN with no buffer. The elution gradient was set as: 0 min, 95% B; 2 min, 95% B; 20 min, 5% B; 23 min, 5% B; 24 min, 95% B; 35 min, 95% B. The column temperature was 35 °C, and flow rate was 0.15 mL/min. The injection volumes were optimized to 1 μL and 6 μL for the urine and saliva samples, respectively. To correct the signal intensities, a series of QC samples were injected with different loading amounts in three technical replicates. For urine samples, the serial QC samples were injected at 0.1, 0.2, 0.4, 0.6, 0.8, 1.0, 1.2, 1.4, 1.6, 1.8, and 2.0 μL (11 volumes in total). For saliva samples, the serial QC samples were injected at 0.6, 1.2, 2.4, 3.6, 4.8, 6.0, 7.2, and 8.4 μL (8 volumes in total). The MS data was collected in negative ion and data-dependent acquisition mode. The MS settings were: capillary voltage, 3.0 kV; nebulizer gas, 1.6 bar; dry gas, 7 L/min; dry gas temperature, 220 °C; mass scan range, 65-1500 (*m/z*); spectra rate, 8.00 Hz; and cycle time, 3.0 s. For centroid spectra calculation, the peak summation width was 3 pts. The mass spectrometer was calibrated using sodium formate.

### Data Processing and Statistical Analysis

The raw LC-MS data was first converted to abf files using Reifycs Abf Converter (ver. 4.0.0). The converted files were processed in MS-DIAL (ver. 4.70) for peak detection, peak alignment, peak area integration, and metabolite annotation.^20^ The detailed settings of MS-DIAL are available in in **SI Text S-1**. Further metabolite annotation was performed on MS-FINDER (ver. 3.52).^21^ The feature annotation results were manually verified using HMDB.^22^

To assess the intragroup variation, pooled relative standard deviation (PRSD) and pooled relative median absolute deviation (PRMAD) were calculated. In brief, when there are *m* biological groups and each group has at least two replicates, we use *Xt* to denote the intensity vector of a feature in group *t* and *SD_Xt_* to denote the standard deviation of intensities in group *t.* The *n_Xt_* represents the number of samples within group *t*. The PRSD was calculated as Eq. 1

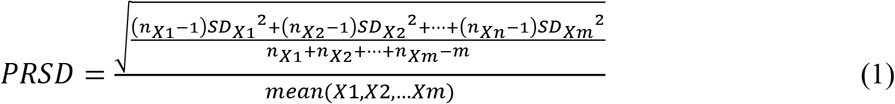

The PRMAD was calculated as Eq. 2

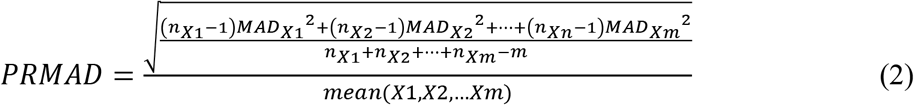

where *MAD_Xt_* denotes the median absolute deviation of intensities in group *t*.

In addition, the relative log abundance (RLA) plot was used to compare the data before and after normalization.^23^ The RLA plot was made using R package *metabolomics* (ver. 0.1.4). Principal component analysis (PCA) was performed on MetaboAnalyst 5.0.^24^ Log-transformation and autoscaling was used to pretreat the data before PCA analysis. To select the significantly altered metabolic features, the *F*-test was first applied to each feature to compare the group variance (unequal variance: *p* < 0.05). Two-sided unpaired Student’s *t* test was performed, and the *F*-test result was used to determine whether equal or unequal variance should be used. The *p* values from the *t* test were further corrected using Benjamini & Hochberg correction method. One-sided Mann–Whitney *U* test was used to compare PRSD and PRMAD between different normalization methods. The *p* value correction and *U* test was performed in R using package *stats* (ver. 4.1.0).

### High-Quality Feature Selection

MAFFIN extracts high-quality metabolic features using the following criteria: (1) their averaged intensities in biological samples should be higher than their intensities in the method blank (default:sample average intensity > 2 × blank intensity); (2) eluted during the analytical gradient; (3) low relative standard deviation (RSDs) of signal intensities in the replicate QC injections (default: RSDs < 25%, according to FDA’s guideline^25^); (4) their MS intensities are highly correlated with their loading amount in serial QC samples (default: Pearson correlation > 0.9). The high-quality metabolic features can be further refined by examining their chromatographic peak shapes and grouping degenerate features, which are features arising from the same metabolite.^26^ In this work, chromatographic peak shapes were manually checked, and degenerate features were grouped using the MS-DIAL curation result (**SI Text S-2**).

### MS Signal Intensity Correction

MS signal intensity correction was performed to convert the MS signal intensity to QC loading amount. In brief, the serial diluted QC samples with different loading amounts are first analyzed alongside biological samples. The feature intensities will be corrected if they have enough data points in serial QC samples and their MS intensities are highly correlated with serial QC loading amounts (Pearson correlation > 0.9). For each feature fulfilling these requirements, a calibration curve was built using the serial diluted QC data, and the signal intensities of the metabolic feature in the biological samples were then converted to QC loading amounts. The R code for data processing was adapted from a previous publication.^27^

### Maximal Density Fold Change

The maximal density FC (MDFC) between two samples is calculated from the FC distribution. First, the FCs of all high-quality features are calculated and log2 transformed. A kernel density function is used to estimate the probability density of the transformed FCs. The kernel density computation was achieved using the kernel “gaussian” in the *stats* R package. In particular, the bandwidth is critical to kernel density calculation, which determines the impact of each data point on the estimated density function. We proposed a method to automatically optimize the bandwidth to ensure normalization accuracy. That method iterates over a range of bandwidths and chooses the bandwidth that achieves the best sample normalization result (detailed in **SI Text S-3**). For the simulated data set where only two samples are involved, the bandwidth was optimized using “SJ” method embedded in the *density()* R function.^28^

### Simulated Data Set

A simulated data set was generated to demonstrate that MDFC can accurately estimate the normalization factor (*k*) between two samples, even when the percentage of up-regulated features is different from that of down-regulated features (i.e., imbalanced data). To initiate the simulation, three parameters were required, including a normalization *k*, percentage of unchanged features, and percentage of up-regulated features (the percentage of down-regulated features can then be calculated). Two samples were simulated with 5,000 features in total. For the unchanged features, their intensities in sample 1 and sample 2 were generated from the same normal distribution. In this case, the intensities of all features in sample 1 (IS_1_) have

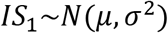

The intensities of unchanged features in sample 2 (IS_2-un_) have

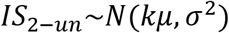

For the up- or down-regulated features, their intensities in sample 1 and sample 2 were generated from two normal distributions with a certain FC of mean values (*f*). Compared to sample 1, the intensities of up- (IS_2-up_) and down- (IS_2-down_) regulated features in sample 2 have

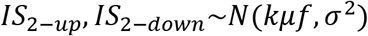

where *f* is higher than 3/2 when up-regulated, and lower than 2/3 when down-regulated. To obtain a comprehensive evaluation, we tested different percentages of unchanged and up-regulated features. The percentage of unchanged features varies from 30 to 100% in steps of 1%. The percentage of up-regulated features varies from 0 to 100% of the unchanged features in steps of 1%.

### Comparison to Other Normalization Methods

MAFFIN workflow was compared to four commonly used post-acquisition sample normalization methods, including sum normalization, median normalization, PQN normalization, and quantile normalization. These four normalization methods were performed on MetaboAnalyst 5.0.^24^

## Results and Discussion

### MAFFIN Workflow

The schematic workflow of MAFFIN is shown in **Figure 1**. To perform MAFFIN, a set of serial diluted QC samples (or QC samples with different loading amounts) need to be analyzed alongside the real metabolomics samples. The input of MAFFIN is a metabolic feature intensity table, which is generated by conventional metabolomics data processing, including metabolic feature extraction and retention time alignment.^20, 29^ Before performing MDFC-based sample normalization, two data preprocessing steps are needed. First, high-quality features were selected because many detected features are not real metabolites or metabolites with poor quantification performance. These low-quality features include but are not limited to features that: 1) are background noise; 2) have low technical reproducibility in LC-MS analysis; 3) have low Pearson correlation between sample loading amount and measured signal intensity. These metabolic features do not represent real metabolites in biological samples. Therefore, their intensity values should not be considered in sample normalization. However, in conventional post-acquisition sample normalization methods, these low-quality features are not excluded from calculating normalization factors, which might cause biased sample normalization. It is important to note that the concept of choosing high-quality metabolic features for sample normalization should also be applicable in nuclear magnetic resonance (NMR)-based metabolomics.

**Figure 1.**
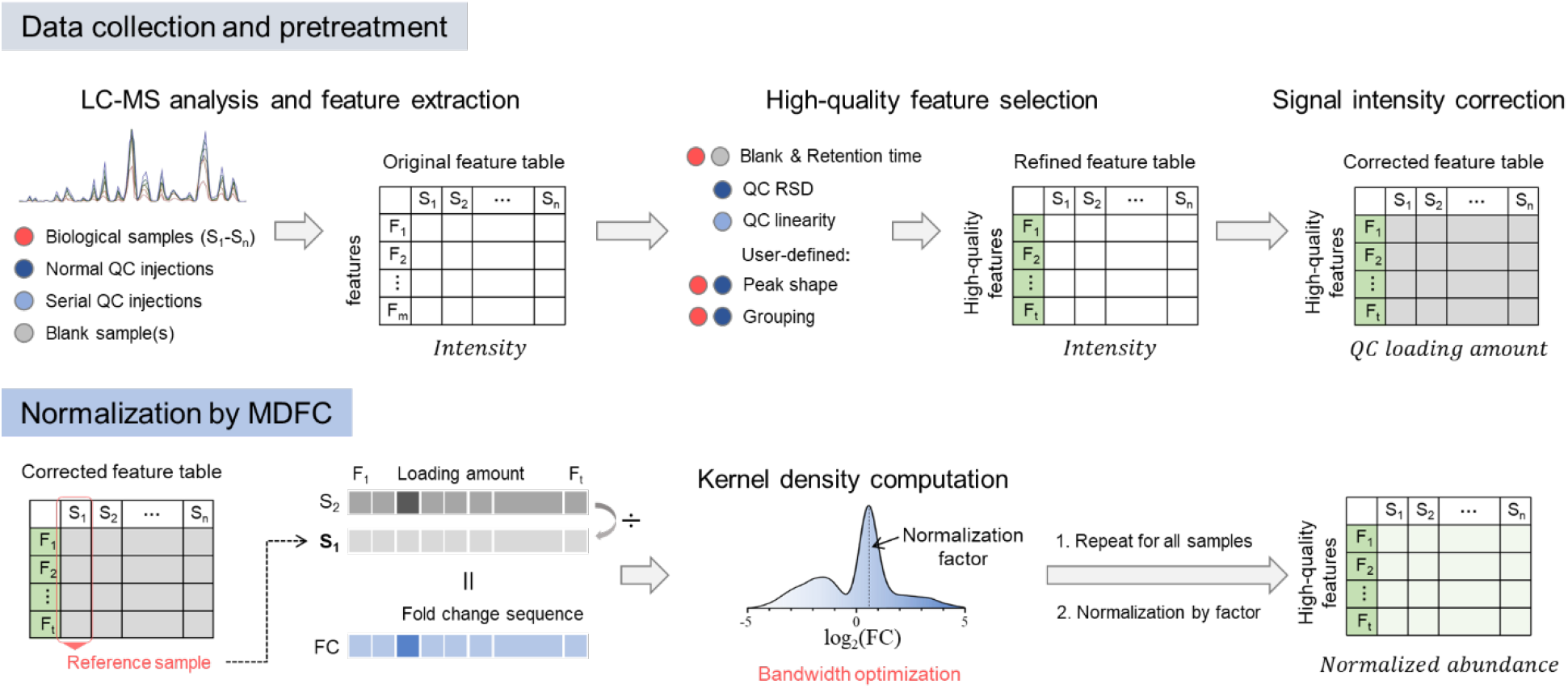
Schematic workflow of MAFFIN.

Another important feature of MAFFIN is to correct MS signal intensities before FC calculation. Our previous works show that the MS signal FCs can be inflated or compressed for most metabolic features even in the linear response range of electrospray ionization.^27, 30^ In MAFFIN, we adapted the previously published MRC workflow to convert MS signals to QC loading amounts.^27^ The FCs calculated using QC loading amounts represent the real concentration FCs and benefit the following estimation of maximal density FC. It is important to note that both high-quality feature extraction and MS signal intensity correction are optional in the MAFFIN workflow, but we highly recommend these two steps, as they can significantly improve the normalization accuracy.

After high-quality metabolic feature selection and MS signal intensity correction, the refined intensity table is used to calculate MDFC for normalization. Its fundamental theory is that for two samples, the unchanged features should have very similar FCs due to proportional total sample amount or concentration differences, while the up- or down-regulated features have different FCs. If we calculate the density of FC distribution of all features, the FC at the highest density (i.e., MDFC) will be caused by unchanged features, thus represents the sample concentration ratio. To calculate MDFC, the sample with the most features detected is first selected as the reference sample. FCs of metabolic feature intensities are then calculated between the test sample and the reference sample. Next, the FCs are log-transformed to make them symmetric around the nonchanged metabolites. After that, Gaussian kernels are used to estimate a density function of the FC distribution, and the FC at the maximal density is used as the normalization factor. Compared to histogram-based discrete estimation, kernel density simulation is continuous and thus has a more accurate estimation of normalization factor.

### Selection of High-Quality Metabolic Features

We first demonstrated the importance of selecting high-quality metabolic features for post-acquisition sample normalization. First, we randomly picked two urine samples from the human urine data set and applied high-quality feature selection. As we can see in **Figure 2A**, the low-quality features have a FC distribution distinct from that of the high-quality features. In particular, the MDFC of low-quality features is 1.34, while the MDFC of high-quality features is 2.05. This demonstrates that if the low-quality features are not excluded, the estimated normalization factor will be biased. Therefore, it is important to select the high-quality metabolic features for post-acquisition sample normalization.

**Figure 2.**
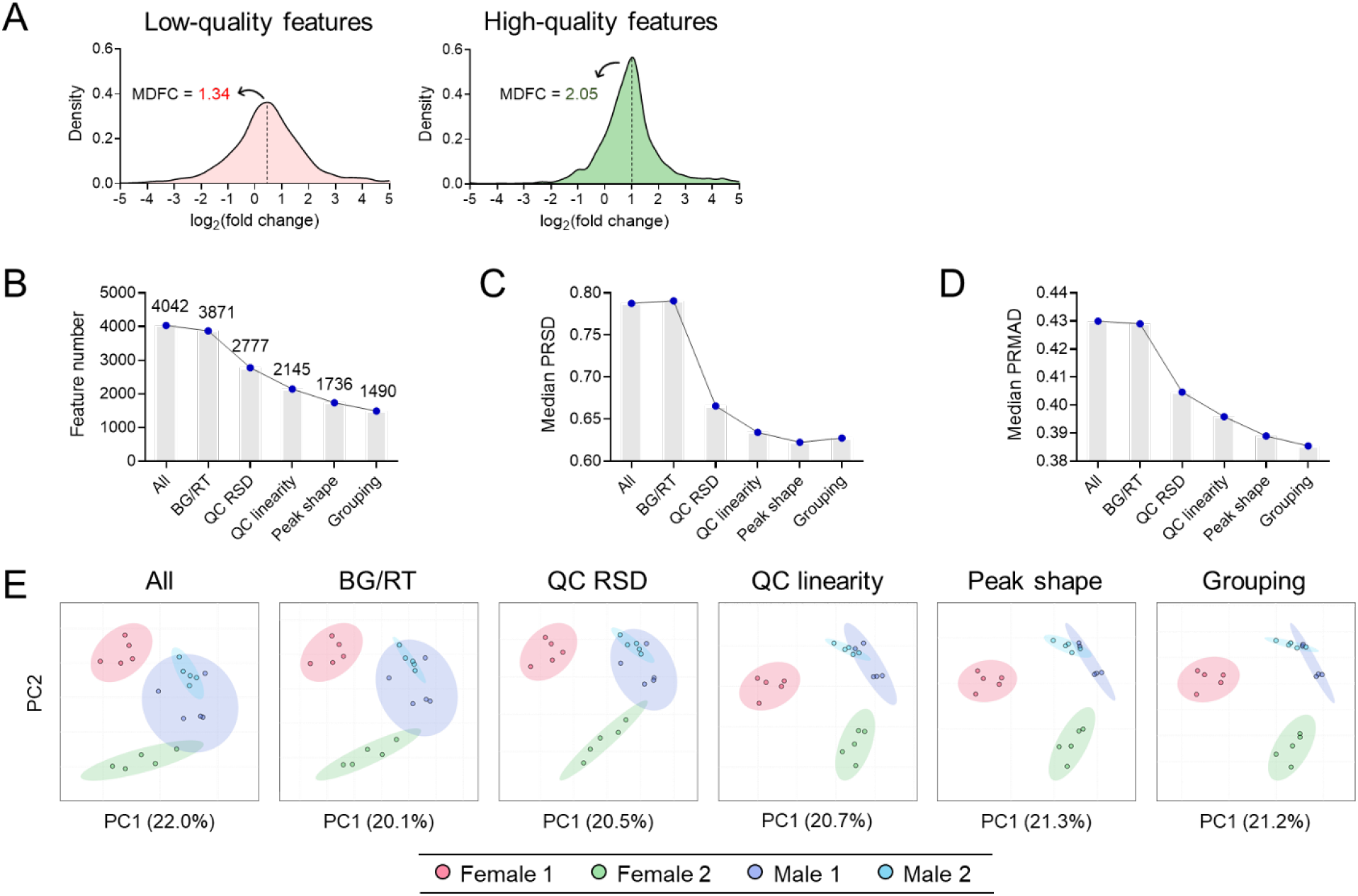
High-quality feature selection in MAFFIN. (A) The FC distributions of low- and high-quality features from two urine samples. (B) Feature number; (C) median of PRSD; (D) median of PRMAD; and (E) PCA score plot after applying different feature selection criteria.

We further evaluated the impact of high-quality feature selection on normalization using the complete urine metabolomics data set. In **Figure 2B**, the metabolic feature number decreased from 4042 to 1490 after all filters were applied. The drop in feature number indicates that the features with good analytical performance might only account for a small portion of all detected features, and we believe that only these features should be selected for sample normalization.^31^ After applying each filter, we corrected the MS signal intensities and normalized the data using MDFC. We then calculated the intragroup variation for each feature using PRSD and PRMAD, as both are measures of the variability of feature intensities. In more specific terms, PRSD uses the square root of the deviation from the mean, while PRMAD uses the median of the absolute deviations from mean. As they capture different data information, we used both to achieve a comprehensive evaluation. Since each feature provides a PRSD and a PRMAD, we examined the median of PRSD and PRMAD of all features and plotted them in **Figures 2C** and **2D**. We observed that the filter of “QC RSD” reduced intragroup variation the most. This suggests that it is critical to remove features with large RSDs in technical replicates prior to normalization factor calculation. Furthermore, the normalization results were evaluated using PCA, as a better normalization method leads to better data separation in the PCA score plot.^32–34^ As shown in **Figure 2E**, a clear trend of improved data separation is observed with more filters applied to select high-quality features.

It is also important to note that the two last steps, selecting features by peak shape and feature grouping, can also help reduce the intragroup variation. For features with bad peak shapes, properly determining their peak areas is a challenge for feature extraction software.^35^ Therefore, features with poor chromatographic peak shapes should be excluded from the calculation of normalization factor. In addition, feature grouping clusters features from the same metabolite and keeps only one feature for normalization, ensuring that all unique features have the same weight for normalization. However, the PRSD results indicate that these two steps do not have a strong effect on variation reduction. Therefore, these two steps are not included in MAFFIN. If needed, users can incorporate other existing bioinformatics tools to select features with good chromatographic peak shapes and/or conduct feature grouping. For example, *peakonly*^36^ or EVA^37^ can be used to remove features with poor chromatographic peak shapes. On the other side, mz.unity,^38^ ISFrag,^39^ or MS-CleanR^40^ can be used for feature grouping.

### MS Signal Correction

Next, we used the serial diluted QC samples in the urine data set to demonstrate that the FCs calculated from signal intensities should not be used directly without signal correction. To demonstrate this, we evaluated whether the metabolic feature FCs are a good estimation of the dilution factor. In principle, the dilution factor should be easily estimated using the feature FC at the maximal count. As shown in the light gray histograms in **Figure 3A,** our results of dilution factors 3, 6, and 10 indicate that the FCs calculated directly from signal intensities might not be a good representation of the true dilution factor. In particular, at higher dilution factor, the maximal count disappears. In comparison, after applying signal intensity correction, the FC at the maximal count can always be easily recognized (the black gray histograms in **Figure 3A).**

**Figure 3.**
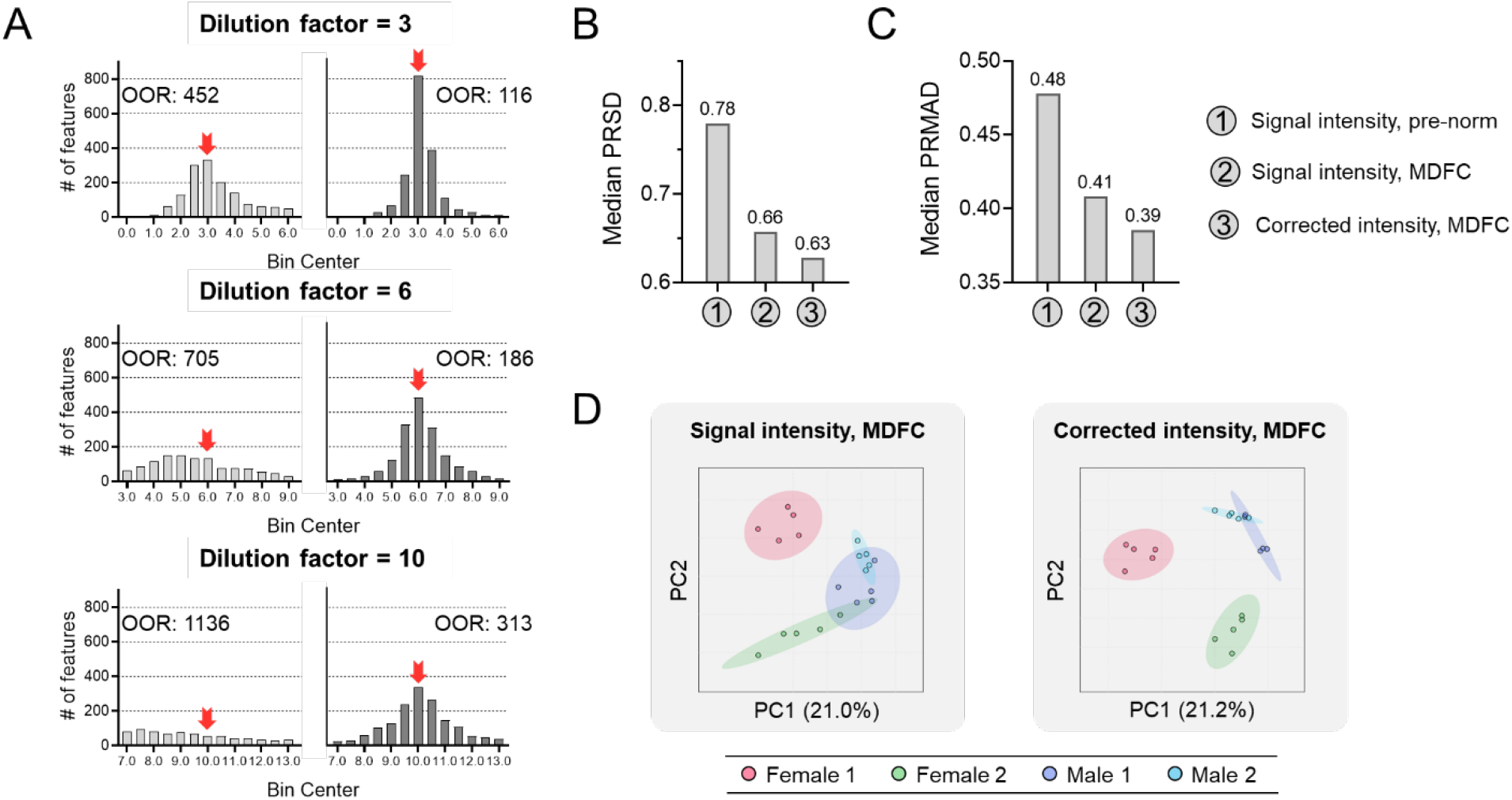
MS signal intensity correction in MAFFIN. (A) The FC distributions of technically diluted urine samples with different dilution factors. The histograms in light and dark gray are from measured MS signal intensity and corrected intensity, respectively. The red arrow indicates the position of the correct dilution factor. OOR: number metabolic features that are out of the plotting range. (B) Median of PRSD; (C) median of PRMAD; and (D) PCA score plot of normalization result before and after intensity correction.

Next, we studied how MS signal intensity correction improves the normalization results. We used the high-quality features selected and corrected their MS intensities according to the MAFFIN workflow. Both corrected and non-corrected data were normalized using the MDFC-based method. We plotted the PRSD and PRMAD of normalized data, including corrected and non-corrected data, as well as the raw intensity data (no normalization applied) results. As shown in **Figures 3C** and **3D,** the intragroup variation was clearly reduced after intensity correction. In addition, a clearer data separation was observed on the PCA score plot (**Figure 3E)**, which confirms the improved data quality after MS signal correction.

### MDFC vs. Median FC

We further compared MDFC with conventionally used median FC to show its advantages in estimating normalization factors. The median FC-based normalization was reported to correct the NMR-based metabolomics data.^18^ Although easy to calculate, median is not a robust estimator of normalization factors. To demonstrate this, we first simulated a two-sample data set of imbalanced percentages of up- and down-regulated features with a true dilution factor of 1.5. In this simulation, we first assumed three types of features, which are unchanged, up-, and down-regulated. We then fixed the percentage of unchanged features (50%) and simulated different percentages of up-regulated features from 5 to 45% in steps of 5%. As shown in **Figure 4A**, MDFC shows a stable and accurate estimation of normalization factor (1.5001 ± 0.003) across the testing percentages. In comparison, the median FC estimated normalization factors ranging from 1.39 to 1.63, when up-regulated features account for 5% and 45%. This demonstrates that median FC has poor normalization factor estimates when the data is imbalanced. We made a more comprehensive evaluation by testing different percentages of unchanged, up-, and down-regulated features at two other true dilution factors (2.0 and 5.0). In each case, the differences between the estimated normalization factors and the true dilution factors were calculated and presented in color gradient. As shown on the left side of **Figure 4B,** when the number of up-regulated features are different from the number of down-regulated features, positive and negative errors were observed in median FC-based method. In comparison, on the right side, the MDFC-based method is robust and never shows any bias across the testing range. Furthermore, comparing the results of different dilution factors, the median FC bias gets bigger (the colored area gets larger) as the true dilution factor increases, which suggests that median FC is less accurate for biological samples of significantly different total sample amount or metabolite concentrations.

**Figure 4.**
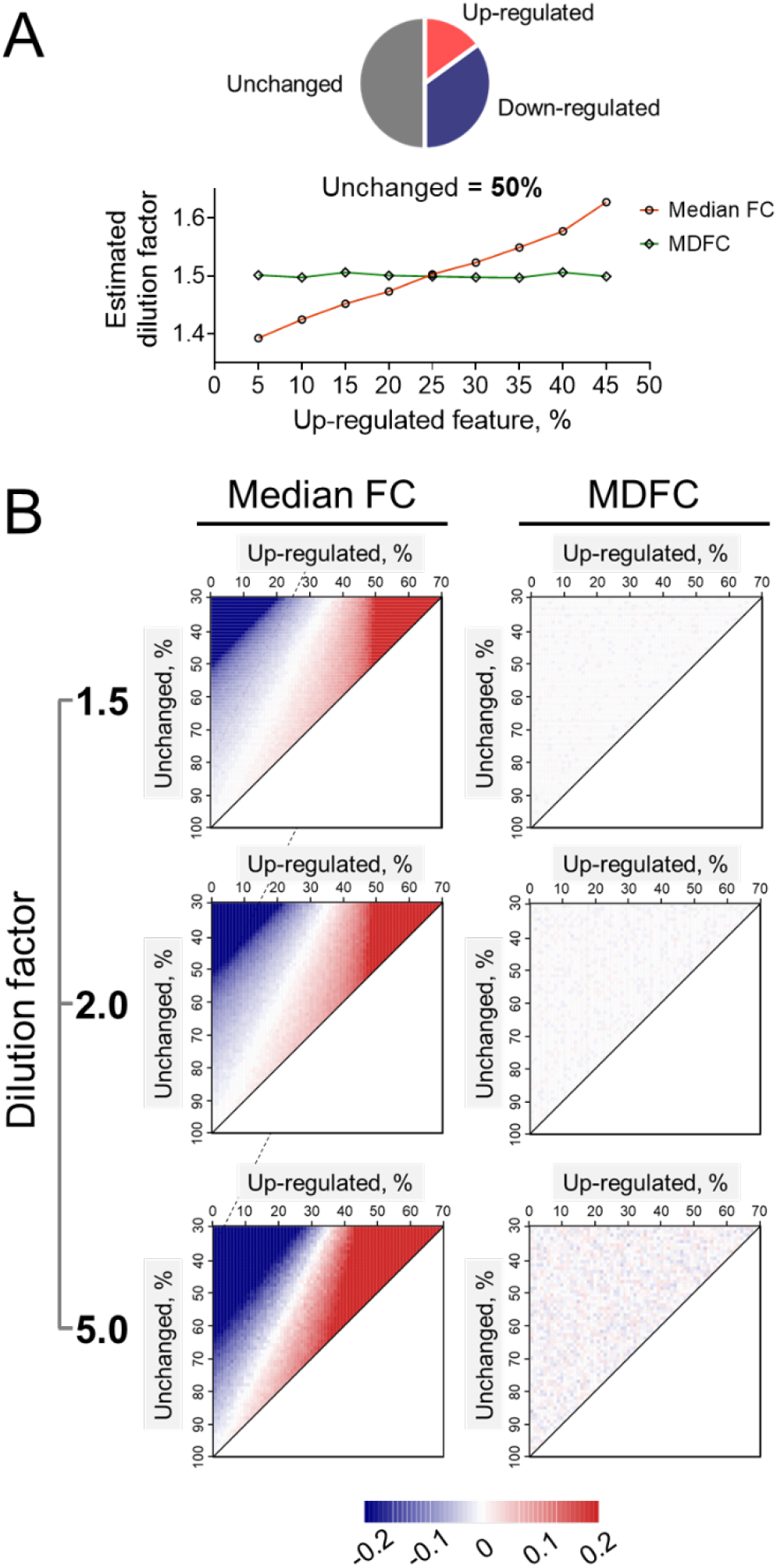
Comparison of MDFC and median FC. (A) Estimation of dilution factors by MDFC and median FC when unchanged features account for 50%. (B) Heatmaps show comprehensive comparison of MDFC and median FC with different data structures.

### Performance of MAFFIN

The performance of the MAFFIN workflow was evaluated and compared to four other post-acquisition normalization methods using the same human urine data set. The RLA plot was first used to visualize the overall sample concentration difference. To make the RLA plot, all feature intensities were log-transformed, and for each feature, its transformed intensities were subtracted by the median value, log transformed. The scaled intensities were visualized using box plot.^23^ Our results show that before sample normalization, a clear biological dilution effect exists in urine samples (**Figure 5A**). Interestingly, after sample normalization by MAFFIN, the urine metabolomics data show little overall concentration variance (**Figure 5B**), indicating that MAFFIN can effectively reduce biological dilution effect.

**Figure 5.**
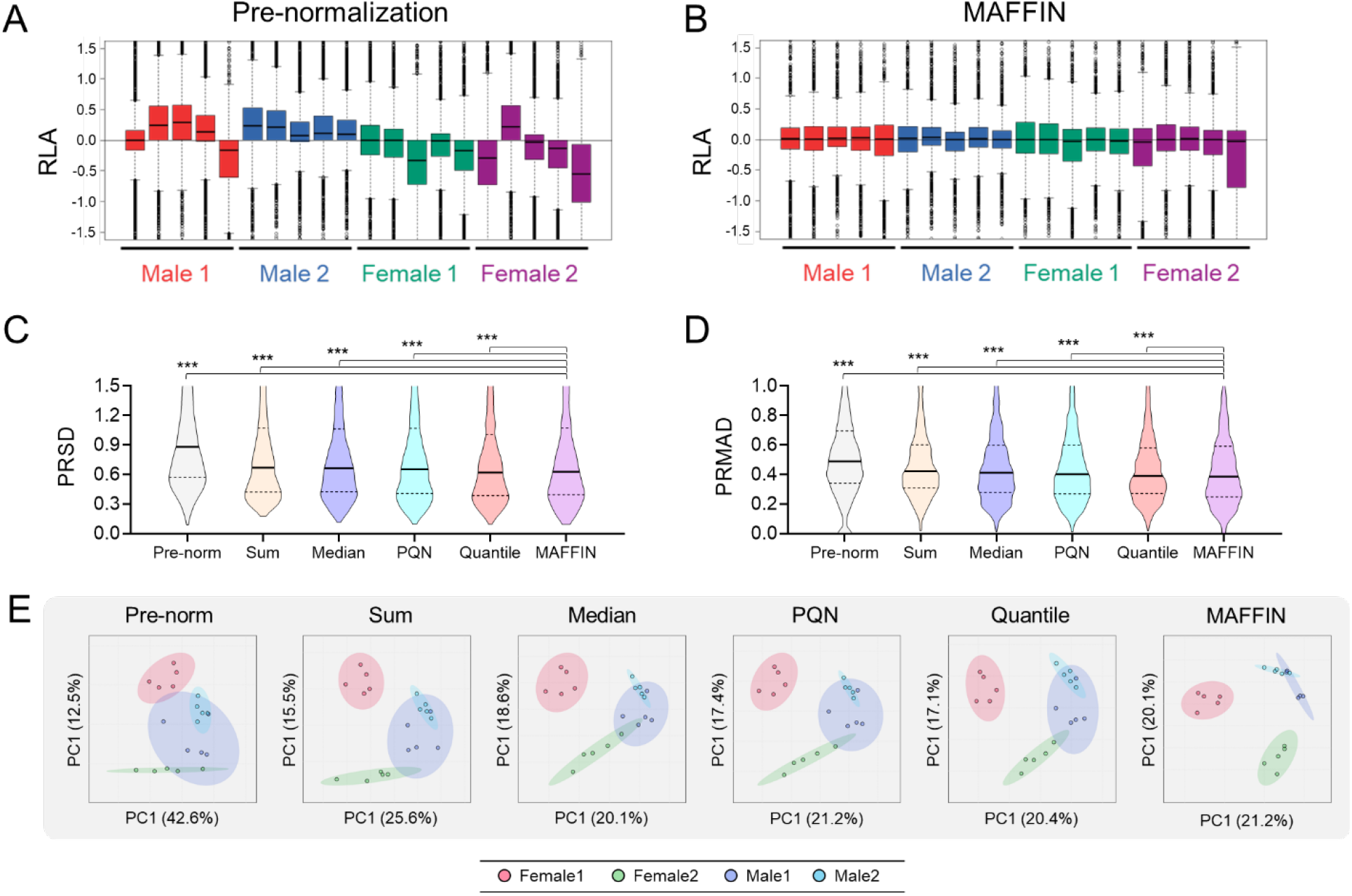
Performance of MAFFIN and comparison to other methods. RLA plots of urine data (A) before and (B) after MAFFIN normalization. Violin plots show the (C) PRSD and (D) PRMAD distributions of different data normalization methods. (E) PCA score plots of data normalized by different methods.

Next, we compared MAFFIN to other normalization methods. In this work, sum, median, PQN, and quantile normalization were compared, all of which are post-acquisition sample normalization methods. After normalizing the urine metabolomics data by the different methods, their intragroup variations, including PRSD and PRMAD, were calculated and plotted in violin plots as shown **Figures 5C** and **5D**, respectively. In both plots, the MAFFIN algorithm provides the lowest intragroup variation. We further performed an one-tailed U test, which examines whether MAFFIN results have statistically lower PRSD or PRMAD than the other methods. As presented in **Figures 5C** and **5D**, the data processed by MAFFIN have significantly lower intragroup variation than the other methods (*p* < 0.001). We further compared MAFFIN to other methods using PCA analysis. As presented in **Figure 5E**, MAFFIN shows the best data separation in PCA score plot. There are many explanations for the lower normalization powers of the commonly used algorithms. The sum normalization result is severely influenced by features with high abundance.^18^ The median normalization assumes that the median signal intensities should be same in different samples. However, the median signal intensity does not mean median metabolite concentration, as different metabolites have different ionization efficiencies. The PQN method uses median FC to estimate the dilution factor, which is biased with imbalanced data. The quantile normalization assumes all data has the same distribution, which is too strict and not in line with reality.^17^ Besides the inherent limitations, the lack of feature selection and MS signal intensity correction in these four methods also contribute to their poor performance. Collectively, these results suggest that MAFFIN can effectively reduce the intragroup variation, serving as a robust and universal normalization method to remove variation.

Finally, MAFFIN was demonstrated on a human saliva metabolomics work composed of 10 male and 10 female saliva samples. LC-MS analysis of these saliva samples detected 2092 metabolic features. Among them, 847 were selected as high-quality metabolic features in MAFFIN (**Figure 6A**), and the sample with the most detected features was used as the reference sample (**Figure 6B**). After MS signal correction, the normalization factors for each sample were determined (**Figure 6B**). It is interesting to see that the reference sample, which has the most detected features, also has the largest normalization factor (factor = 1, **Figure 6B**). This is reasonable as samples with more metabolic features are usually more concentrated. From the normalization results, we found that the normalization factors have an RSD of 24.2%, and the highest sample concentration was 2.72-fold of the lowest one. These results suggest there is a large dilution effect in the saliva samples and that sample normalization is necessary to remove the intrasample variance.

**Figure 6.**
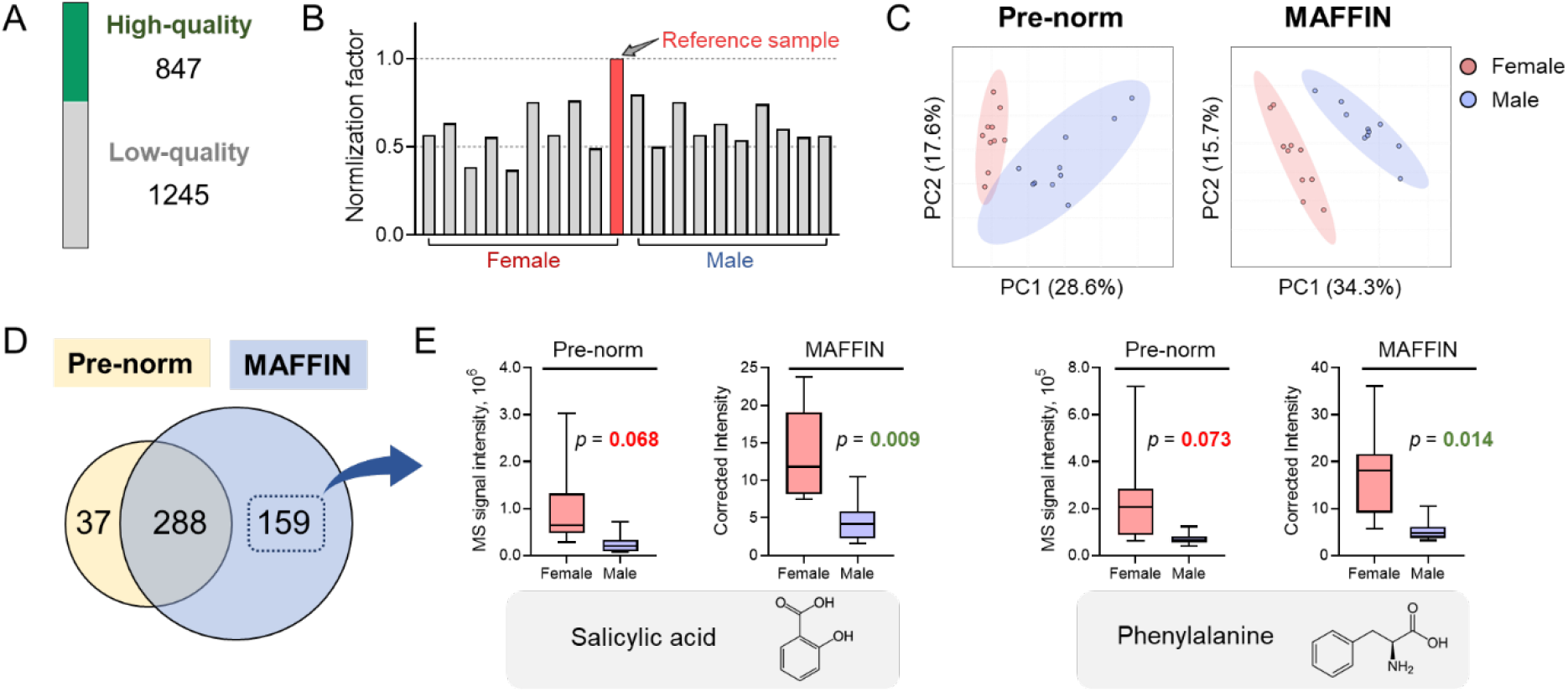
Biological application of MAFFIN on human saliva samples. (A) The high-quality feature selection result. (B) The normalization factor calculation results. (C) PCA results before and after MAFFIN normalization. (D) Venn diagram shows the difference of significant metabolites before and after normalization (adjusted *p* < 0.05, FC < 0.667 or > 1.5). (E) Box plots show two newly discovered significant metabolites, salicylic acid and phenylalanine.

The benefit of MAFFIN can be clearly seen, as post-normalization separation of two groups on a PCA score plot is more distinct (**Figure 6C**). Furthermore, the differential metabolic analysis was applied on the data before and after normalization, and *t* test was utilized to select the significantly altered metabolic features (adjusted *p* < 0.05, FC < 0.667 or > 1.5). Among the 2092 total features, there are 159 newly discovered significant features after applying the MAFFIN algorithm, while 37 features became insignificant after normalization (**Figure 6D**). We further manually annotated all the newly discovered significant features by matching their experimental MS^2^ spectra to publicly available databases^20^ and in-silico MS^2^ databases^21^. The annotations were also manually verified using HMDB.^22^ The feature annotations and *t* test results are detailed in **SI Table S-1**. As shown in **Figure 6E**, we presented the concentration levels of two annotated metabolites, which became significant after MAFFIN-based sample normalization. Among them, salicylic acid was reported to appear in human saliva and used in clinical pharmacokinetic studies.^41, 42^ In addition, phenylalanine has also been confirmed to be critical to oral health.^43, 44^ In summary, MAFFIN is a robust normalization method to help reveal metabolome differences in case studies with high accuracy and confidence.

## Conclusion

In this work, we propose MAFFIN, a novel sample normalization strategy that normalizes untargeted metabolomics data collected from MS-based platforms. Our results show that by incorporating high-quality feature selection, MS signal intensity correction, and MDFC, MAFFIN outperforms commonly used post-acquisition sample normalization strategies in terms of reducing intragroup variation and improving separation in PCA plots. MAFFIN was also applied on a saliva metabolomics study and identified 159 additional significantly altered metabolic features related to sex difference. As a post-acquisition normalization method, MAFFIN requires no extra measurements on samples and shows accurate and robust normalization results for both balanced and imbalanced data. MAFFIN can be extremely useful for biological samples that lack a quantity that reflects the total sample amount or concentrations of metabolites. These include saliva, sweat, fecal content, and many others. Furthermore, MAFFIN can also be adapted for non-biological samples such as waste water or other environmental samples used for exposome research. We believe that the development of MAFFIN addresses the critical challenge of sample normalization in untargeted metabolomics. It advances the standardization of untargeted metabolomics workflows and broadens metabolomics applications in biomarker discovery, mechanistic understanding, and beyond.

## Supplementary Information

**Table S-1.** Annotation and *t* test results of newly discovered significant metabolic features; **Text S-1.** Parameter settings in MS-DIAL; **Text S-2.** Feature grouping based on MS-DIAL curation results; Text S-3. Bandwidth optimization.

## Acknowledgment

This study was funded by University of British Columbia Start-up Grant (F18-03001), Canada Foundation for Innovation (CFI 38159), New Frontiers in Research Fund/Exploration (NFRFE-2019-00789), National Science and Engineering Research Council (NSERC) Discovery Grant (RGPIN-2020-04895). We also thank Alisa Hui for proof reading this manuscript.

