## Supplementary Information for "MAFFIN: Metabolomics Sample Normalization Using Maximal Density Fold Change with High-Quality Metabolic Features and Corrected Signal Intensities"

Internet: https://huan.chem.ubc.ca/

### Outline

**Table S-1**. Annotation and *t* test results of newly discovered significant metabolic features.

**Text S-1.** Parameter settings in MS-DIAL.
**Text S-2.** Feature grouping based on MS-DIAL curation results.
**Text S-3.** Bandwidth optimization.

**Table S-1**. Annotation and *t* test results of newly discovered significant metabolites.

| Metabolite name | p before MAFFIN | p after MAFFIN | Quality |
|---|---|---|---|
| 12-HETE | 0.0810 | 0.0312 | High-quality |
| 13-HODE | 0.0550 | 0.0252 | High-quality |
| 2-Hydroxy-4-methylpentanoic acid | 0.0732 | 0.0252 | Low-quality |
| 2-Pyrocatechuic acid | 0.0722 | 0.0224 | High-quality |
| Adenosine monophosphate | 0.0604 | 0.0187 | High-quality |
| Cascaroside C | 0.0630 | 0.0497 | High-quality |
| Cellobiose | 0.1394 | 0.0403 | High-quality |
| Citrulline | 0.1332 | 0.0188 | High-quality |
| Di-2-furanylmethane | 0.0698 | 0.0119 | High-quality |
| Ethyl 2-furanacrylate | 0.0644 | 0.0284 | High-quality |
| Fucose | 0.1950 | 0.0277 | High-quality |
| Glucosamine 6-phosphate | 0.0836 | 0.0337 | Low-quality |
| Hydrocinnamic acid | 0.0512 | 0.0349 | Low-quality |
| Inosine | 0.0592 | 0.0342 | High-quality |
| Leucic acid | 0.0677 | 0.0094 | Low-quality |
| LPE 18:1 | 0.0731 | 0.0188 | Low-quality |
| LPE 18:2 | 0.0589 | 0.0252 | Low-quality |
| LPI 18:0 | 0.0749 | 0.0284 | Low-quality |
| Mannose 1-phosphate | 0.0733 | 0.0197 | High-quality |
| PE (20:3/16:0) | 0.0512 | 0.0130 | Low-quality |
| PG (16:0/18:1) | 0.0671 | 0.0175 | High-quality |
| PG (16:0/20:4) | 0.0986 | 0.0398 | High-quality |
| Phenol | 0.0661 | 0.0278 | Low-quality |
| Phenylalanine | 0.0731 | 0.0142 | High-quality |
| Phosphoethanolamine | 0.0540 | 0.0271 | High-quality |
| Phosphoglyceric acid | 0.0519 | 0.0492 | Low-quality |
| Phosphoric acid | 0.0479 | 0.0181 | High-quality |
| PI (20:4/18:0) | 0.0540 | 0.0176 | Low-quality |
| Prehumulinic acid | 0.0540 | 0.0111 | High-quality |
| PS (18:0/18:2) | 0.0734 | 0.0201 | High-quality |
| PS (18:0/20:0) | 0.0574 | 0.0189 | Low-quality |
| PS (22:0/18:1) | 0.0792 | 0.0234 | Low-quality |
| Quinic acid | 0.0986 | 0.0218 | High-quality |
| Salicylic acid | 0.0677 | 0.0093 | High-quality |
| Theophylline | 0.0937 | 0.0347 | High-quality |
| Thymine | 0.0524 | 0.0222 | Low-quality |
| Tyrosol | 0.0520 | 0.0392 | High-quality |
| UDP-glucose | 0.0585 | 0.0058 | High-quality |

**Text S-1.** Parameter settings in MS-DIAL.

**Mass accuracy:** $MS^1$ tolerance: 0.01, $MS^2$ tolerance: 0.05

**Peak detection:** Minimum peak height: 1000 amplitude; mass slice width: 0.05 Da

**Identification:** MSMS library: http://prime.psc.riken.jp/compms/msdial/main.html

Accurate mass tolerance ($MS^1$): 0.01

Accurate mass tolerance ($MS^2$): 0.05

The retention time is not used for scoring.

**Adduct:** $[M-H]^-$, $[M-H_2O-H]^-$, $[2M-H]^-$

**Alignment:** Retention time tolerance: 0.3 min; $MS^1$ tolerance: 0.01 Da

**Text S-2.** Feature grouping based on MS-DIAL curation results.

The curation result from MS-DIAL reported the relationship between different features. For example, "found in higher mz's MsMs" means a feature is found in another feature's MS2 spectrum, which indicating this feature is a potential in-source fragment of another feature. We first grouped the features using the curation results. For the grouped features, we further decided if they are different species from the same metabolite. Our hypothesis is that if two features are from the same metabolite, they should have high correlation among samples. Therefore, we calculated the Pearson correlation between pairs of features within a group. The hierarchical clustering analysis (HCL) was used to further cluster the features within a group based on their Pearson correlations. The result of HCL is a tree, and the *cutree()* function (R package *stats*) was used to further separate the features into more groups. A cluster dendrogram is shown below, and the total 10 features were further cut into 5 data groups.

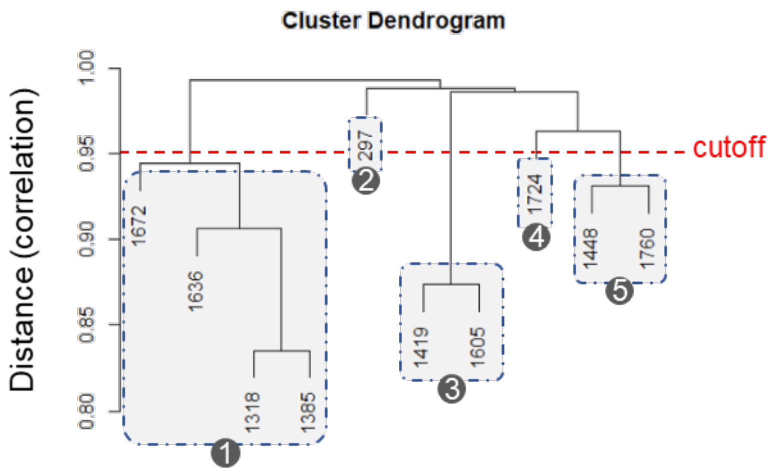


For the features in a data group, we have clear evidence that they are from the same metabolic feature. Therefore, only the feature with the highest average intensity would be kept as a unique metabolic feature.

**Text S-3.** Bandwidth optimization.

The function *density()* computes kernel density estimates, and bandwidth is the most important parameter in this function. Properly choosing bandwidth is vital for an accurate normalization. Here, we present a new bandwidth optimization method for metabolomics data normalization. The rationale is that the best bandwidth achieves the best normalization result, which means the lowest intragroup variation. The intragroup variation was determined by the pooled relative median absolute deviation (PRMAD). To optimize the bandwidth, we computed the PRMAD of all features from a given data set using different bandwidths from 0.1 to 10 (step = 0.1). The best bandwidth achieves the lowest median of PRMAD.
